# AI-Driven Toolset for IPF and Aging Research Associates Lung Fibrosis with Accelerated Aging

**DOI:** 10.1101/2025.01.09.632065

**Authors:** Fedor Galkin, Shan Chen, Alex Aliper, Alex Zhavoronkov, Feng Ren

## Abstract

Idiopathic pulmonary fibrosis (IPF) is a condition predominantly affecting the elderly and leading to a decline in lung function. Our study investigates the aging-related mechanisms in IPF using artificial intelligence (AI) approaches. We developed a pathway-aware proteomic aging clock using UK Biobank data and applied it alongside a specialized version of Precious3GPT (ipf-P3GPT) to demonstrate an AI-driven mode of IPF research. The aging clock shows great performance in cross-validation (R2=0.84) and its utility is validated in an independent dataset to show that severe cases of COVID-19 are associated with an increased aging rate. Computational analysis using ipf-P3GPT revealed distinct but overlapping molecular signatures between aging and IPF, suggesting that IPF represents a dysregulation rather than mere acceleration of normal aging processes. Our findings establish novel connections between aging biology and IPF pathogenesis while demonstrating the potential of AI-guided approaches in therapeutic development for age-related diseases.

## Introduction

Idiopathic pulmonary fibrosis (IPF) is a chronic, progressive lung disease characterized by the excessive accumulation of extracellular matrix components, leading to declining lung function and ultimately respiratory failure. Predominantly affecting individuals over the age of 60, the correlation between aging and IPF underscores the importance of understanding the aging-related mechanisms contributing to its pathogenesis. Both aging and fibrotic diseases pose major healthcare challenges. As such, global deaths to fibrotic diseases have been on the rise over the last three decades and constitute 18%, according to the latest estimates (1). Similarly, the population aging problem has been a mainstay of biomedical and political debates for decades (2,3). Identifying the mechanisms shared by these aging and fibrosis is crucial for developing targeted therapies that can potentially benefit the global population.

Current treatments for IPF are limited and primarily focus on slowing progression rather than addressing underlying causes. Lung transplantation remains the only way to improve a patient’s survival rate, and prior anti-fibrotic therapies may be considered a way to help a patient outlast the long waiting period (4,5). This scarcity of effective therapies stems partly from an incomplete understanding of the molecular and cellular processes driving fibrosis in the aging lung. However, novel strategies to fight IPF are emerging with many of them focusing on its aging-related nature (6).

Recent advancements in biomedical research, notably in artificial intelligence (AI), offer a new vector to developing IPF therapies. AI-driven approaches can analyze vast amounts of biological data to identify novel biomarkers, therapeutic targets, and actionable insights. The existing AI pipelines have been successfully used to analyze the aging footprints in proteomic, transcriptomic, epigenetic and other types of omics biodata (7–9).

Today, these research technologies have matured enough to be used in more practical applications as testified by a number of granted deep aging clock patents (10–13). Even more AI applications are actively used biomedical research settings not focused on aging, such as target discovery, drug candidate design, clinical trial design, and others (14–16). These AI models serve to open new classes of drugs with promising anti-aging potential, such as HIF-PH inhibitors for the treatment of IBD (17,18). Models such as those from the Precious and Nach0 lineups hold the promise of enabling a fully digital mode of clinical research by emulating real-life experimental settings based on existing data (19–22). Simultaneously, LLM-inspired genomic AI models allow scientists to discover new mechanistic and evolutionary theories of aging and explain the behavior of living systems in environments previously considered too complex to model (23–27).

The rapid pace of AI innovation and hardware improvements promotes an optimistic outlook on the future of clinical therapeutics. By extracting actionable knowledge from a diverse and vast range of biomedical studies, more connections between seemingly disparate pathological processes can be built to find the cures for conditions beyond the reach of contemporary medicine. IPF is such a disease with an unclear etiology and a paucity of life-extending clinical countermeasures, apart from lung transplantation (28). The existing body of evidence suggests that the onset of IPF has a strong an aging-related component, suggesting that this disease may be a specific instance of the general aging process (29). Various authors highlight that IPF is characterized by the accumulation of senescent cells, insufficient autophagy, proinflammatory environment, and mTOR deregulation, which are commonly considered hallmarks of aging (30,31). Other authors suggest that the activation of embryonic pathways is essential for lung regeneration, while their deregulation is common in older individuals which leads to aberrant tissue maintenance (32).

In this study, we aim to explore the similarities between aging and IPF using AI models, such as Precious3GPT (P3GPT) and aging clocks. By identifying age-related biomarkers and therapeutic targets and utilizing AI to predict disease status, we offer new avenues for developing novel anti-fibrotic treatments. This study represents a significant step forward in understanding and addressing the complexities of IPF and aging mechanisms with the potential to improve outcomes for patients suffering from this challenging condition.

## Materials and Methods

### Data collection

For aging clock training, we used the UK Biobank collection of 55319 proteomic Olink NPX profiles annotated with age and gender. The validation set was obtained from (33) and the corresponding Dryad repository (34).

For training ipf-P3GPT, we used a filtered collection of prompts used in training the original P3GPT model that contains 3873 prompts, including 672 prompts with the disease2diff2disease instruction and 3201 with the age_group2diff2age_group instruction (**Supplementary File 1**). The disease2diff2disease prompts were further enriched to include differentially expressed genes from *in vivo* and *in vitro* datasets available via Gene Expression Omnibus (GEO). The included studies feature a variety of settings involving fibrotic diseases and their models, such as IPF, liver cirrhosis, NASH, alcoholic liver disease, chronic kidney disease, TGF-β cell models, keloid scarring (**Supplementary File 2**). The full list of the added studies and their differentially expressed features obtained from them are available in the supplementary files. Differential gene analysis for the added transcriptomic studies was carried out using Limma (35,36).

All the prompts were modified to contain only the gene names represented in the Olink 3072 platform (37). The prompts were filtered to keep only those with >100 significantly differently expressed genes (absolute log2 fold change > 1, q-value<0.05).

All prompts were further extended with XML-like tags denoting EFO identifiers of samples’ tissue and its hierarchical ancestors, EFO identifiers of the condition studied in a dataset and its hierarchical ancestors, as well as the EFO identifier of the tissue primarily affected by the condition of interest. Terms from EFO release version 3.73.0 were used for the mapping of GEO-specified metadata to prompt tags. Additionally, we extended each prompt with a binary <is_fibrosis> tag that serves as a direct way to assess whether an observed omics signature is associated with fibrotic changes.

### Proteomic aging clock

We developed a multi-task neural network architecture that combines biological age prediction with pathway activity inference (**Figure 1**). The network consists of three main components: a shared feature extraction backbone, an age prediction branch, and a pathway prediction branch modulated by an attention mechanism.

**Figure 1.**
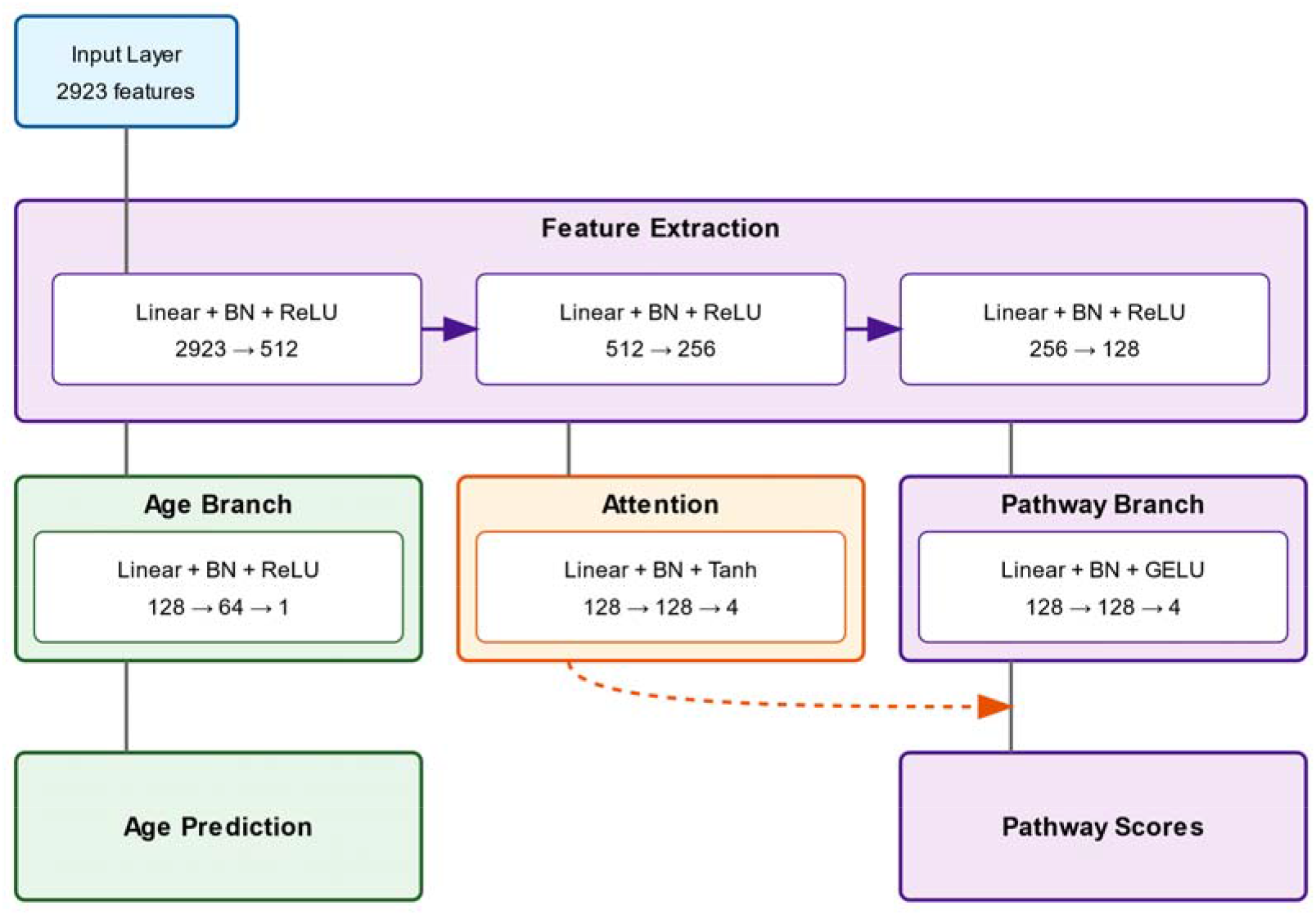
Pathway-aware proteomic aging clock architecture. The neural network consists of a shared feature extraction backbone processing 2923 protein measurements, an age prediction branch, and a pathway prediction branch with attention mechanisms. The model combines age prediction with pathway activity inference for TGF-β signaling, ECM remodeling, inflammation, and oxidative stress, using attention weights initialized from RNA study evidence.

The shared feature extraction backbone processes 2923 protein measurements through three fully connected layers (2923→512→256→128) with batch normalization and ReLU activation. This shared representation feeds into both the age prediction and pathway branches. The age prediction branch consists of two fully connected layers (128→64→1) with batch normalization and ReLU activation, followed by a softplus activation and scale-shift layer to constrain outputs to a biologically plausible age range.

The pathway branch processes the shared features through a fully connected layer with batch normalization and GELU activation (128→128), followed by a final linear layer with tanh activation (128→4) to predict pathway scores. An attention mechanism, implemented as two fully connected layers (128→128→4) with batch normalization and tanh-softmax activation, modulates the pathway predictions.

The attention mechanism is initialized using weights derived from RNA study evidence described in the previous section. These initial weights are calculated by combining differential expression data across multiple studies, with each study weighted based on tissue relevance, disease type, and comparison setting. The attention weights can be updated during training while maintaining biological plausibility through a KL-divergence regularization term.

The model is trained using a combined loss function that incorporates age prediction error, pathway prediction error, and attention regularization. The age prediction component uses relative error to account for age-dependent uncertainty. The pathway prediction component uses mean squared error between predicted and directly calculated pathway scores. The attention regularization term uses KL-divergence to maintain consistency with the biologically-derived initial weights while allowing for data-driven adaptation.

Training is performed using the Adam optimizer with cosine annealing learning rate scheduling and early stopping based on validation loss. Gradient clipping is applied to ensure stable training. The model implementation uses PyTorch and includes batch normalization and dropout (p=0.2) in the feature extraction and age prediction branches to prevent overfitting. At inference, the missing protein values imputed with sample means. See **Supplementary File 3** for the package used to train the clock.

### ipf-P3GPT training

We trained ipf-P3GPT, a lightweight transformer model, to learn differential gene expression patterns in pulmonary fibrosis. The model architecture comprises a 64-dimensional embedding layer, three transformer blocks with two attention heads each, and a vocabulary-sized output layer (**Figure 2, Supplementary File 4**). The model was implemented in PyTorch 2.0.

**Figure 2.**
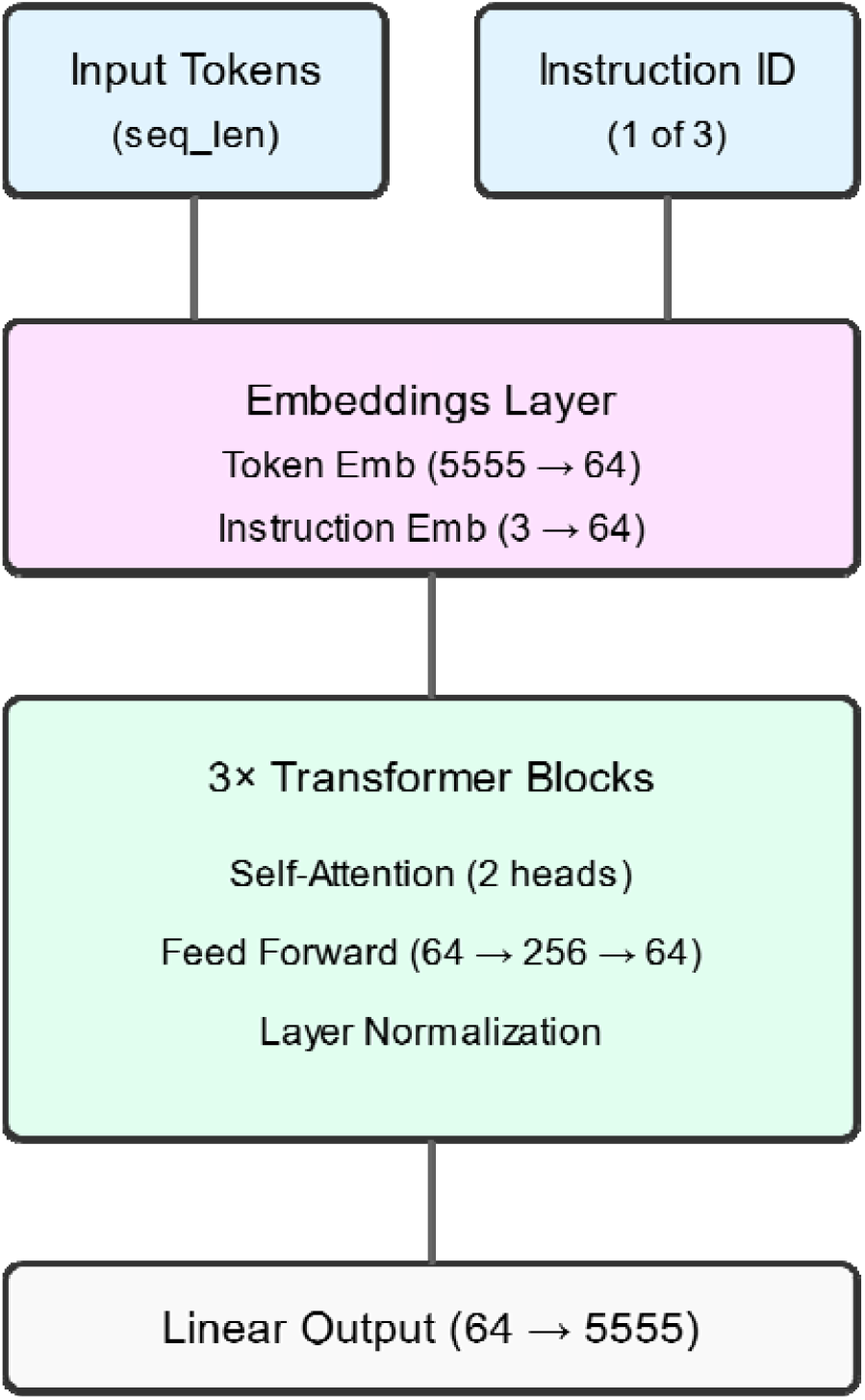
ipf-P3GPT model architecture. The transformer model features a 64-dimensional embedding layer, three transformer blocks with dual attention heads, and a vocabulary-sized output layer. The model processes disease, age group, and compound treatment comparisons using a custom XML-aware tokenizer, achieving 72.2-75.9% validation accuracy across instruction types.

The transformer blocks consist of multi-head self-attention layers followed by feed-forward networks (FFN). Each FFN expands the 64-dimensional input to 256 dimensions through a linear transformation, applies GELU activation, and projects back to 64 dimensions. Layer normalization is applied after both attention and FFN components. The model processes three instruction types: disease comparisons, age group comparisons, and compound treatment comparisons.

Training utilized a custom XML-aware tokenizer with a vocabulary size of 5,555 tokens. Input sequences were padded or truncated to 512 tokens. We employed mixed-precision training with gradient scaling using an AdamW optimizer (learning rate=1e-4) and batch size of 32. The training data was split 90:10 for training and validation, with weighted random sampling to balance instruction types.

The model was trained for 100 epochs, achieving a final validation loss of 2.03. Instruction-specific accuracies reached 72.2% for age group comparisons and 75.9% for disease comparisons. Model weights were saved at the best validation loss checkpoint.

All hyperparameters and architectural choices were empirically determined through ablation studies, with the final configuration optimizing for both model performance and computational efficiency.

### Statistical tests

Treatment effects were analyzed using Mann-Whitney’s U tests to compare healthy versus afflicted groups. For the linear effects assessment in the validation COVID-19 dataset, the Ordinary Least Squares module from the statsmodels for Python package.

### Pathway analysis

We analyzed four key molecular pathways relevant to IPF pathogenesis: TGF-β signaling, ECM remodeling, inflammation, and oxidative stress. For each patient, pathway scores were calculated as aggregated standardized values of proteins known to participate in respective pathways, with protein-pathway assignments based on curated hallmark sets from MSigDB (38–40).

The biological aging clock used in this analysis was trained using a pathway-aware architecture that incorporated prior knowledge of molecular mechanisms involved in IPF and aging. This was achieved by including pathway-specific attention layers in the neural network, allowing the model to learn pathway-specific contributions to biological age predictions. The pathway attention weights were initialized using a curated database of aging-related pathway signatures and were further refined during model training on the UK Biobank proteomic dataset.

### Manuscript Preparation

The initial manuscript draft was created using DORA (Draft Outline Research Assistant, https://dora.insilico.com/), an AI-powered scientific writing platform. DORA is Insilico Medicine’s LLM-based assistant for automated scientific writing, leveraging an ensemble of over 50 specialized AI agents powered by large language models. These agents work in concert to gather relevant literature, analyze data, and generate high-quality scientific content. All agents are empowered with Retrieval-Augmented Generation (RAG) technology, which enables comprehensive data collection while maintaining scientific accuracy via online fact-checking and PubMed citation linking. After DORA generated the initial draft, all authors collaborated to critically review, expand, and refine the manuscript, ensuring scientific rigor and the accuracy of the presented statement and references.

## Results

### Fibrosis-aware aging clock

We first developed a proteomic aging clock using data from 55,319 UK Biobank participants aged 50-85 years (mean age 69 years). The clock achieved a mean absolute error (MAE) of 2.68 years and R^2^=0.84 in five-fold cross-validation (CV) when using the full list of 2923 available features, demonstrating robust age prediction capabilities (**Figure 3A, Supplementary File 5**). After confirming the clock’s accuracy in CV, we applied it to the Olink Explore 1536 dataset from (33) featuring healthy, moderate, and severe cases of COVID-19 (N=84). Due to many missing values in this dataset and non-uniform age distribution across outcome groups, a direct comparison of prediction errors between them was not feasible. Hence, we applied a linear regression method to assess the effect of disease severity on the pace of aging. Our analysis identified that the patient with a sever case of the infection, and thus likely to develop lung fibrosis, had significantly higher biological age compared to healthy controls (+2.77 years, p=0.026), suggesting that the trained clock carries biological relevance in fibrotic cases (**Supplementary File 6**).

**Figure 3.**
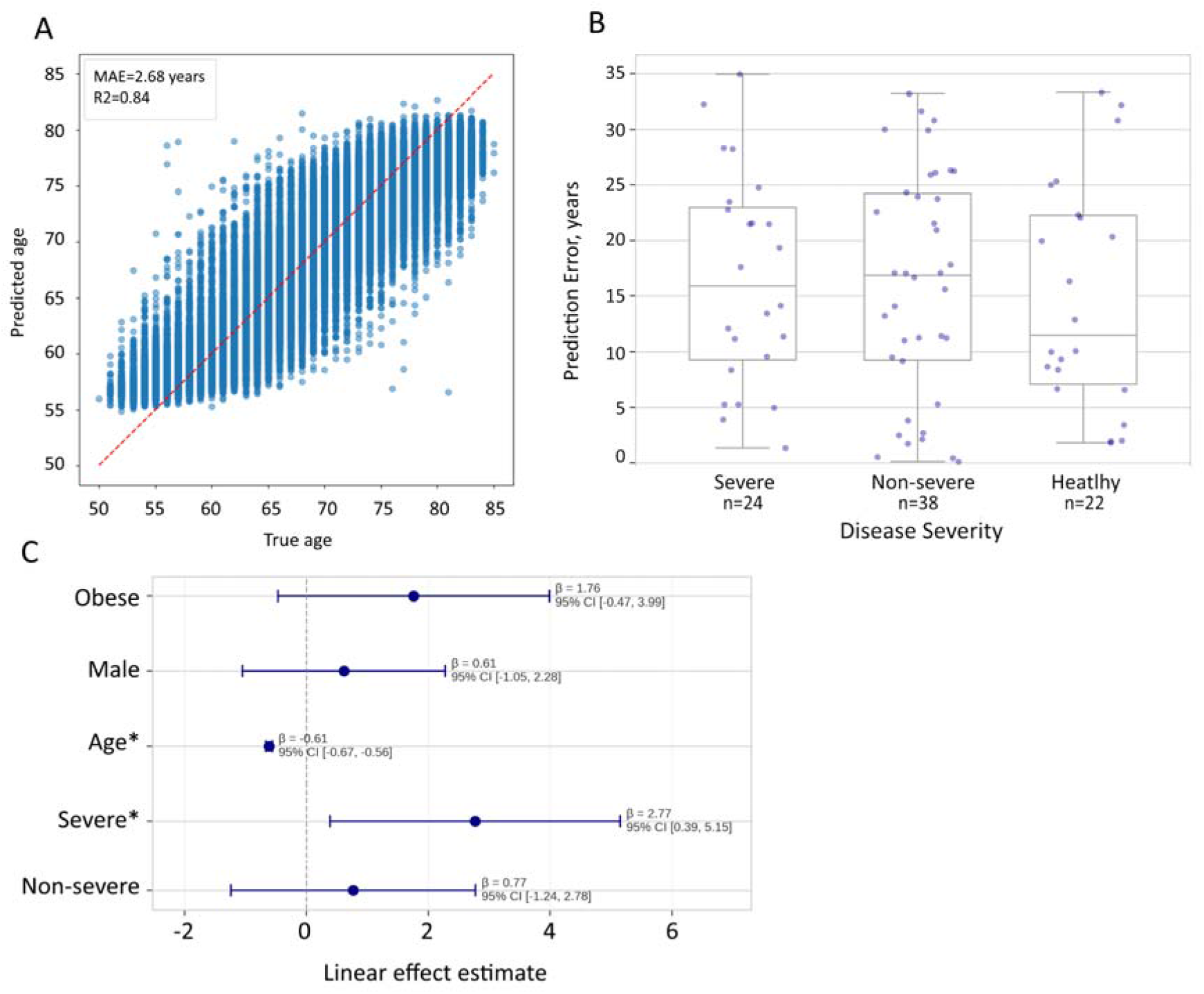
Comparison of biological age predictions between healthy controls and cases of severe COVID-19 infection. (A) Proteomic aging clock shows R2=0.84 in the task of age prediction in CV within the UK Biobank dataset (N = 55,319). (B) Biological age acceleration (difference between predicted and chronological age) across severity groups. compared to healthy controls. Error bars represent standard error of the mean. (C) Linear regression analysis reveals that patients with severe cases, which are likely to develop lung fibrosis, showed significantly higher biological age predictions (+2.77 years, p=0.026).

### ipf-Precious3GPT analysis

After exploring the function of the aging clock, we set out to identify the genes contributing to the progression of both IPF and aging by using ipf-P3GPT. ipf-P3GPT is an abridged version of the full-scale P3GPT that was trained on a data collection enriched in fibrotic disease human and in vitro studies. Using this model, we generated two distinct gene expression profiles: one representing the classical IPF transcriptomic response and another modeling the aging process in lung tissue from 30 to 70 years old. The model assigned attention scores to each gene, indicating their relative importance in the respective biological processes.

Analysis of differentially expressed genes revealed distinct transcriptional signatures for IPF (n=96 genes) and aging-associated (n=93 genes) processes in lung tissue (**Table 1, Supplementary File 7**). The overlap between these signatures was limited to 15 genes (15.6% of IPF signature), suggesting substantial divergence in the underlying molecular programs. Among the overlapping genes, 46.7% (7/15) showed concordant directional changes, while 53.3% (8/15) exhibited opposing regulation between IPF and aging conditions. This initial observation prompted us to investigate the specific molecular pathways affected in each condition.

**Table 1A.** Shared Genes Between IPF and Aging-Associated Lung Fibrosis.

| Gene | IPF |  | Aging |  |
| --- | --- | --- | --- | --- |
|  | Direction | Score | Direction | Score |
| **Concordant genes** |  |  |  |  |
| IL1RL1 | ↑ | 48.8 | ↑ | 16.9 |
| ASPN | ↑ | 43.9 | ↑ | 9.4 |
| PAPPA | ↑ | 34.7 | ↑ | 12.5 |
| F2R | ↓ | 34.4 | ↓ | 5.6 |
| COL15A1 | ↑ | 32.3 | ↑ | 3.4 |
| ADAMTS1 | ↓ | 23.3 | ↓ | 2.4 |
| IL6 | ↑ | 19.2 | ↑ | 3.6 |
| CD83 | ↓ | 16.1 | ↓ | 3.3 |
| **Discordant genes** |  |  |  |  |
| COL1A1 | ↓ | 51.3 | ↑ | 10.0 |
| SERPINA3 | ↓ | 43.2 | ↑ | 9.2 |
| FOSB | ↑ | 41.3 | ↓ | 7.7 |
| EDNRB | ↓ | 37.9 | ↑ | 14.2 |
| APOH | ↓ | 36.1 | ↑ | 9.4 |
| NPTX2 | ↑ | 27.5 | ↓ | 9.2 |
| **GPC5** | ↓ | 24.8 | ↑ | 5.0 |
| **VIPR1** | ↓ | 23.4 | ↑ | 12.5 |
| **QSOX1** | ↑ | 21.9 | ↓ | 4.3 |

**Table 1B.** Key Unique Genes in Each Condition (Top 15 by Attention Score)

| IPF-specific |  |  | Aging-specific |  |  |
| --- | --- | --- | --- | --- | --- |
| Gene | Direction | Attention | Gene | Direction | Attention |
| AKT3 | ↓ | 100.0 | IL10RA | ↑ | 100.0 |
| TUBB3 | ↑ | 88.3 | MFAP4 | ↓ | 61.7 |
| CFB | ↓ | 86.0 | FGF7 | ↓ | 57.4 |
| PKN3 | ↓ | 70.7 | S100A3 | ↑ | 49.6 |
| CCL8 | ↓ | 63.5 | SEMA3G | ↓ | 45.7 |
| GDF15 | ↓ | 60.3 | IL11 | ↓ | 36.1 |
| APOC1 | ↑ | 56.0 | FASLG | ↑ | 30.4 |
| MET | ↑ | 55.6 | NTRK2 | ↑ | 25.5 |
| BCHE | ↓ | 55.6 | VIT | ↑ | 23.2 |
| SFRP1 | ↑ | 52.7 | MMP3 | ↑ | 20.4 |
| AGBL2 | ↓ | 52.5 | SCGB3A1 | ↑ | 19.5 |
| DAPK2 | ↓ | 51.1 | MARCO | ↓ | 19.5 |
| MRC1 | ↓ | 50.7 | TAGLN3 | ↑ | 18.1 |
| NRGN | ↓ | 49.7 | GZMA | ↓ | 17.4 |
| VEGFA | ↑ | 47.9 | FNDC1 | ↑ | 15.6 |

Values represent relative attention scores among the gene tokens attending to the fibrosis diagnosis by the ipf-P3GPT model. ↑ indicates upregulation, ↓ indicates downregulation.

Both conditions demonstrated significant involvement of ECM-associated genes, albeit with distinct regulatory patterns. IPF signature included multiple collagen types (COL1A1↓, COL3A1↓, COL5A1↑, COL15A1↑) and matrix-modifying enzymes (MMP1↑, MMP13↑). The aging signature, in contrast, showed a different matrix remodeling profile, characterized by changes in structural proteins (ELN↓, MFAP4↓, MFAP5↓) and matrix-associated factors (POSTN↓). Notably, COL1A1 showed opposing regulation between IPF (downregulated) and aging (upregulated), suggesting divergent matrix reorganization mechanisms.

Further examination revealed that the IPF signature was enriched for TGF-β pathway components (TGFBR1↓, GDF15↓, BGN↑) and inflammatory mediators (CXCL8↑, IL1B↓, CCL8↓). While the aging signature shared some inflammatory mediators (IL6↑, IL1RL1↑), it exhibited a distinct growth factor profile (FGF7↓, CSF3↑). Both conditions showed involvement of matrix-associated growth factors, though through different molecular effectors.

## Discussion

Our study presents several significant findings regarding the intersection of aging biology and IPF progression, while introducing novel AI-driven approaches for understanding disease mechanisms and therapeutic interventions. To further the community’s understanding of IPF and other fibrotic diseases, we present two deep learning models: a proteomic aging clock developed with UKB data and an abridged version of P3GPT, a transformer-based model for generating and analyzing omics-level data.

The proteomic clock we developed shows an accuracy (R^2^=0.84) below another recently published aging clock ProtAge (R^2^=0.94) in a different subsection of UKB (41). While ProtAge uses fewer features and higher accuracy, we argue that the presented aging clock represents a methodological advancement in biological age assessment. Unfortunately, a direct comparison was not feasible since the authors had not deposited the weights of the model used. For our purposes, the processing of all available proteomic information was essential to incorporate the IPF context in the learned representations. The addition of pathway-level information provides additional biological interpretability and context-specific relevance that distinguishes it from general-purpose aging clocks. The four hallmark pathways added to the model’s attention block and secondary output head were selected based on their reported importance in IPF development (4,42–44). Yet, in future iterations we shall consider including more pathways in the model to expand its field of applications.

We then inspected an Olink proteomic dataset using this clock to find out if its prediction error carries biological signal. Olink platforms are rapidly gaining popularity and we have located multiple open access datasets generated with them, such as those featured in (45–49). These datasets, however, lack sufficient chronological age annotation, describe non-fibrotic diseases, or were generated with Olink platforms measuring <1000 protein quantities. The datasets generated with the most complete Olink Explore 3072 platform remain rare and require a lengthy access procedure. Thus, we focused on the COVID-19 dataset from (33) as the most fitting for our purposes. While not directly representing IPF, the COVID-19 infection has been reported to lead to pulmonary fibrosis in severe cases and thus may be considered biologically relevant for the assessment of our new model (50). The application of the aging clock to this dataset revealed that more severe cases of these respiratory diseases are associated with significantly (p<0.05) higher age predictions. We anticipate that this trend shall be also observed and clearer in other pulmonary and fibrotic diseases.

To further investigate the similarities shared by aging and IPF, we used generative AI in the form of ipf-P3GPT which was instructed to generate transcriptomic lung IPF and aging (from 30 to 80 years) signatures. Both these signatures were identified by the model as the cases of fibrosis, and both contained genes from the key IPF-related pathways highlighted above (ECM remodeling, inflammatory signaling, TGF-β pathways). Yet, at a gene level, the two cases were quite dissimilar with only eight genes showing concordant expression in the emulated processes. Such genes include known contributors to IPF, such as COL15A1 (51). Some known drivers of fibrosis are only present in the IPF signature: MMP1, MMP13, AKT3, IL6 (52–55). And some key ECM components, such as COL1A1 are reported upregulated in IPF and down-regulated in noremal aging, as also recorded in literature (56).

The differential regulation of key fibrotic pathways between IPF and aging generations suggests that IPF may represent dysregulation rather than mere acceleration of normal aging processes. The shared pathway involvement but divergent gene regulation indicates that IPF might arise when normal age-related tissue maintenance mechanisms become pathologically altered. This hypothesis is supported by the opposing regulation of critical ECM components and the partial overlap in inflammatory mediators, suggesting that IPF may hijack normal aging-associated, compensatory tissue remodeling programs, driving them toward a pathological state. Even among genes whose expression, according to ipf-P3GPT, is concordant in aging and IPF, the attention scores assigned by the model vary, indicating a different level of involvement in the accompanying fibrotic processes.

The insights provided by ipf-P3GPT demonstrates the value of creating focused, application-specific AI models. While maintaining core capabilities of the original P3GPT architecture, this streamlined version offers reduced computational overhead, rapid inference times, and domain-specific knowledge concentration. These characteristics make it particularly suitable for exploring disease mechanisms and therapeutic responses in the context of IPF. An important extension to the previously reported P3GPT omics data representation is the addition of a fibrosis-specific tag to all prompts in training, which would allow ipf-P3GPT to be used directly as a tool for assessing the clinical significance of external omics signatures or its own generations. By exploring the gene tokens attending to the fibrotic status, we were able to identify key contributors to the aging and IPF progression.

Future studies incorporating tissue biopsies, rather than blood proteomes, and continuous follow-up would be valuable for validating our findings. As for the AI arm of experiments, a wider set of generations need to be explored, including those representing other fibrotic diseases, to identify more reliable patterns of multi-omic expression.

Our research project opens several promising avenues for investigation. We incentivize other scientists to use the data and models demonstrated here to gain a deeper understanding of the aging and fibrotic processes in their own studies. The particular use cases for the presented models may include indication expansion and the development of novel therapies for age-related diseases. We are also planning to expand our approach to train an array of specialized small-scale P3GPT-like models, each acting as a knowledge source for their respective diseases. Such small-scale AI models can be used to streamline drug development for rare and dangerous conditions and augment existing AI workflows already in use.

The successful application of both our aging clock and ipf-P3GPT demonstrates the growing importance of AI tools in therapeutic development. These approaches not only enhance our understanding of disease mechanisms but also provide frameworks for identifying new therapeutic opportunities (19,20). The pathway-aware architecture of our aging clock in particular represents a step toward more biologically informed AI models that could accelerate the development of targeted geroprotective interventions.

## Conclusion

This study underscores the value of integrating aging biology into therapeutic development for age-related diseases. The combination of targeted therapeutic intervention with AI-driven analysis provides a powerful approach for understanding disease mechanisms and identifying effective treatments. While further research is needed to fully characterize the research utility of ipf-P3GPT and the proteomic clock, our findings suggest they are promising new tools for the study of IPF and potentially other age-related fibrotic conditions.

## Authors contribution

AA — Supervision, Methodology, Writing (review and editing)

AZ — Conceptualization, Funding acquisition, Writing (review and editing)

FG — Data curation, Investigation, Software, Visualization, Writing (review and editing)

FR — Supervision, Methodology, Writing (review and editing)

SC —Formal analysis, Investigation, Writing (review and editing)

## Acknowledgments

The authors thank A.Pogorelskaya for her expertise and assistance in preparing ipfP3GPT training data.

## Data availability

The code and data used in this study are available as an OSF repository (57).

